# Measuring single-virus fusion kinetics using an assay for nucleic acid exposure

**DOI:** 10.1101/2022.05.20.492704

**Authors:** Ana M. Villamil Giraldo, Steinar Mannsverk, Peter M. Kasson

## Abstract

The kinetics by which individual enveloped viruses fuse with membranes provide an important window into viral entry mechanisms. We have developed a real-time assay using fluorescent probes for single-virus genome exposure than can report on stages of viral entry including or subsequent to fusion pore formation and prior to viral genome trafficking. Encapsulating such fluorescence probes in the lumen of target membranes permits specific detection of fusion events and exclusion of leakage without fusion. Using this assay, we show that influenza virus fuses with liposomes of different sizes with indistinguishable kinetics, suggesting that the starting curvature of such liposomes does not control the rate-limiting steps in influenza entry.

## Introduction

Enveloped viruses infect cells via a process of membrane fusion between the viral membrane and a cellular membrane, mediated by viral fusion proteins. Activation of the fusion proteins is often rate-limiting, but at least two free energy barriers and corresponding rate-limiting steps have been identified leading to the lipidic intermediates in fusion ^1-4^. These are the formation of an initial fusion stalk between the proximal leaflets of the viral and cellular membranes and the formation of a fusion pore, at which point the viral and cellular membranes are topologically continuous. Viral genome release and the potential for replication occurs subsequent to fusion pore formation. In some cases, this may be immediate, while in others downstream events have been identified controlling genome release^5, 6^.

Single-particle measurements of viral fusion kinetics have proven highly informative in constraining possible fusion mechanisms^7-17^. Although bulk kinetics have helped identify the timescales for fusion and many of the factors affecting rate-limiting steps^2, 18-27^, the distribution of event waiting times observed in single-particle measurements reports on the degree of functional heterogeneity in the viral population and on which kinetic mechanisms could account for the observed behavior. Analyses in this regard have leveraged substantial prior advances in single-turnover enzymology, which has developed theory and metrics for analyzing such events ^8, 28-30^.

Most single-particle measurements of viral fusion kinetics utilize optical probes: fluorescently-labeled lipids that dequench upon mixing with an unlabeled target membrane^7, 9, 10, 13, 15, 16, 31^, water-soluble dyes loaded at a quenching concentration ^7, 32^, or genetically encoded proteins that change their fluorescent state in a pH-sensitive manner^12, 33, 34^. Aptamer-based detection offers a powerful means to measure viral genomes^35, 36^ but is more challenging to employ for real-time single-virus detection. Subsequent trafficking of viral genomes has also been measured via quantum-dot and other labeling methods^6, 37^. For technical reasons, the use of soluble dyes for content mixing has been more challenging, including unstable loading of the dyes into influenza virions^32^, thermally-induced fusion reported in non-viral systems^38^, and challenges in separating fusion from leakage events ^32^. An additional factor is that the diffusion of different soluble factors through nascent fusion pores may happen at different stages of the process, particularly when comparing small molecules and large proteins ^39-41^. Nevertheless, these experiments have yielded substantial insight into the factors controlling fusion pore formation.

Here, we report the development and use of a small-molecule dye assay for genome exposure, schematized in Figure 1. When this dye is encapsulated in a target liposome, genome exposure could occur either via transit of the viral genome through the fusion pore or (more likely) diffusion of the dye into the viral interior and binding to exposed genomic material. Here, we measure the genome exposure kinetics of influenza virus. Influenza contains eight viral RNA segments, each encapsulated in the virion as viral ribonucleoproteins (vRNPs). A matrix layer (M1) further protects the genome but disassembles in response to viral pH drop when the M2 proton channels in the viral membrane are activated^42-46^. We demonstrate that nucleic-acid-binding dye can efficiently detect influenza genomes and that this occurs relatively rapidly after fusion pore formation and in the absence of other cellular factors. We also test the dependence of viral genome exposure on target liposome curvature. In accordance with our prior results on influenza lipid mixing^47^ but in contrast to prior results on synaptic vesicle fusion^48^, we find that target membrane curvature has no detectable effect on the kinetics of genome exposure or fusion pore formation.

**Figure 1.**
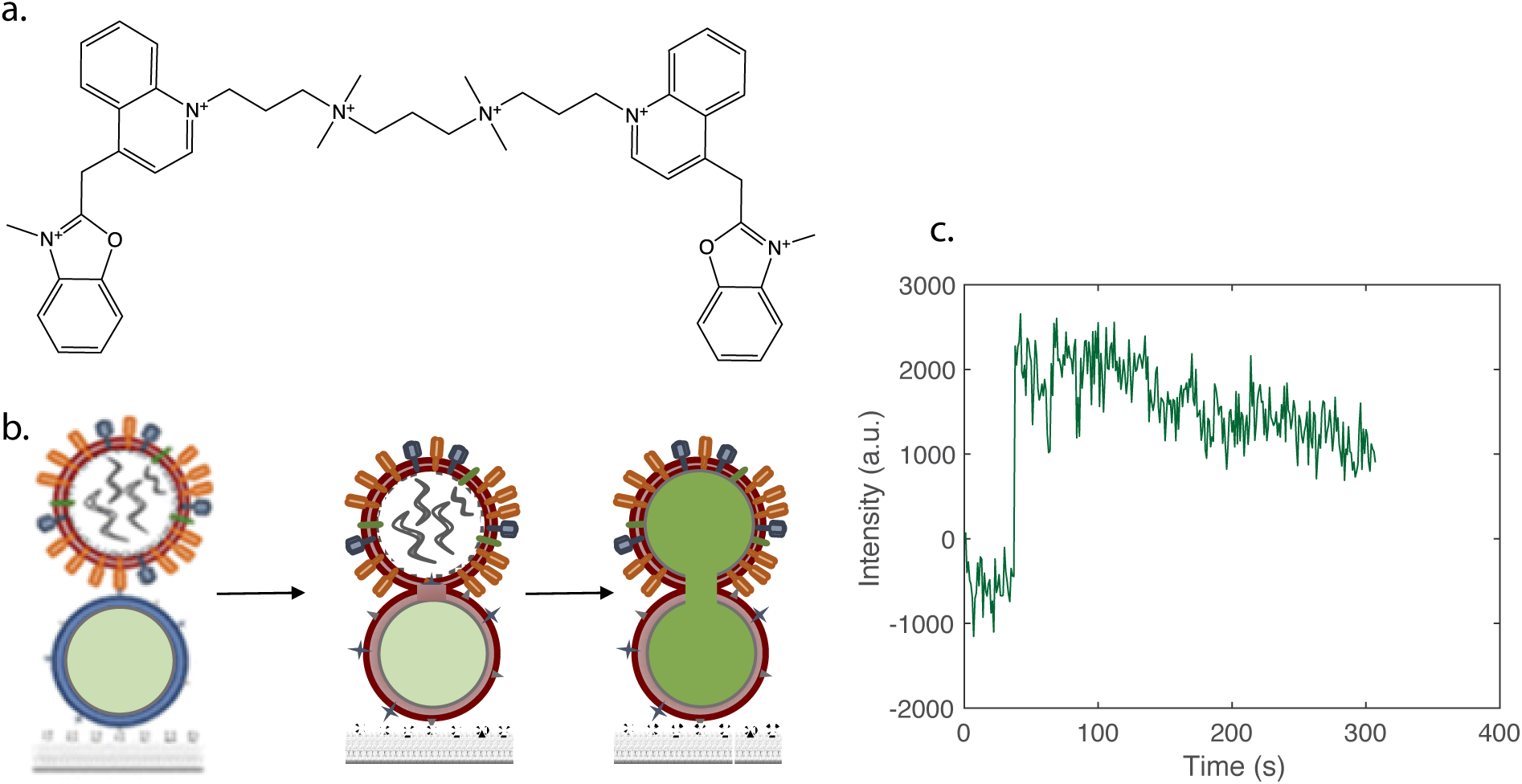
Detection of viral nucleic acid exposure in membrane fusion. DiYO-1 (an oxazole yellow homodimer, panel a) is a nucleic-acid-binding dye that dramatically increases fluorescence quantum yield upon binding to DNA or RNA. Encapsulation of DiYO in liposomes creates a substrate for fusion that is essentially non-fluorescent until after a fusion pore has been opened and the viral genome is accessible for dye binding, schematized in panel (b). A sample fluorescence trace is plotted in panel (c), where the abrupt jump in fluorescence represents an influenza virus fusion and genome exposure event.

## Materials and Methods

### Materials

Palmitoyloleoylphophatidylcholine (POPC) dioleoylphosphatidylethanolamine (DOPE), cholesterol (chol), and biotinylated 1,2-dipalmitoyl-sn-glycero-3-phosphoethanolamine (biotin-DPPE) were purchased from Avanti Polar Lipids. Disialoganglioside GD1a from bovine brain (Cer-Glc-Gal(NeuAc)-GalNAc-Gal-NeuAc) was purchased from Sigma-Aldrich. Texas Red-1,2-dihexadecanoyl-sn-glycero-3-phosphoethanloamine (TR-DHPE) and Oregon Green-1,2-dihexadecanoyl-sn-glycero-3-phosphoethanolamine (OG-DHPE) were purchased from Thermo Fisher. PLL-PEG and PLL-PEG-biotin were purchased from SuSoS AG. DiYO-1 (CAS 143413-85-8) was purchased from AAT Bioquest. X-31 influenza virus (A/Aichi/68, H3N2) was purchased from Charles River Laboratories.

### Liposome preparation

Lipids (67 mol% POPC, 20% DOPE, 10 % chol, 1% biotin-DPPE and 2% GD1a) were mixed in chloroform, which was evaporated under nitrogen to a thin film. Residual chloroform was removed by keeping the samples in vacuum for at least 3 h. The lipid films were hydrated in 10 µM DiYO or 16 mM calcein in 10 mM NaH_2_PO_4_, 90 mM sodium citrate, 150 mM NaCl, pH 7.45. The lipid suspension at a final lipid concentration of 0.56 mM was subjected to 15 freeze-thaw cycles, and large unilamellar vesicles were obtained by extruding the suspension through polycarbonate membrane filters (Avestin) with a nominal diameter of 100 nm or 200 nm. Excess unincorporated dye was removed using a Sephadex G-25 desalting column (Cytiva).

### Viral labeling

6 µl of TR-DHPE (0.75 mg/mL in ethanol) was mixed with 20 mM HEPES, 150 mM NaCl pH 7.2. A small volume of virus suspension (typically 9 μL) at 2 mg total protein/mL was mixed with 4× volume (typically 36 μL) of dye:buffer mixture and incubated for 2 h at room temperature on a rocker. Labeled virus was purified from unincorporated dye by adding 1.3 mL 20 mM HEPES, 150 mM NaCl pH 7.2 and centrifuging for 50 min at 21,000 rcf. The pellet containing labeled virus was resuspended in 10 μL of 20 mM HEPES, 150 mM NaCl pH 7.2 and allowed to rest on ice at 4°C overnight. Labeled virus was either stored on ice at 4°C and used directly or subjected to a single freeze-thaw cycle prior to use. Procedures involving influneza virus were performed under BSL-2 conditions.

### Single-virus fusion assays

Fusion assays were performed as previously described^47^, with the addition of new content-mixing and genome exposure fluorescent probes. Briefly, glass slides were treated with a mixture of 95% PLL-PEG for passivation and 5% PLL-PEG-biotin for functionalization and then incubated for 15 minutes with streptavidin at 0.2 mg/mL. Biotinylated liposomes were then tethered to the surface by incubation overnight at 4°C. Texas-Red-labeled influenza virus was added prior to the start of fusion experiments. At each step, unbound material was washed away using >2 mL of phosphate buffer, pH 7.45. Fusion was initiated via a buffer exchange to pH 5.0 (10 mM NaH_2_PO_4_, 90 mM sodium citrate, 150 mM NaCl).

### Fluorescence microscopy

Micrographs and videos were acquired using a Zeiss Axio Observer inverted microscope and a 100X oil immersion objective. Additional optical parameters included excitation/emission filters (Chroma Technology) at 480/40 and 535/50 for calcein and DiYO and 560/40 and 630/75 for Texas Red and an Arduino controlled Spectra-X LED Light Engine (Lumencor) as a light source. Images were recorded with a Zyla 4.2 sCMOS camera (Andor Technologies) controlled via MicroManager. *Image analysis*. Images were analyzed using previously reported single-virus detection and spot-tracking protocols^15, 49^. MATLAB (The Mathworks) code is available from https://github.com/kassonlab/micrograph-spot-analysis.

### Bulk fluorescence measurements

Lyophilized Poly-A RNA (Qiagen) was resuspended at 1 µg/µL in RNAase-free water with 0.04% sodium azide and then diluted in phosphate buffer (10 mM NaH_2_PO_4_, 90 mM sodium citrate and 150 mM NaCl) at pH 5.0 to reach the designated concentration. The diluted RNA was aliquoted in a 96-well plate, followed by addition of DiYO-1 dye diluted in phosphate buffer, yielding a final dye concentration of 10µM. Fluorescence was measured using a Clariostar Plus plate reader (BMG Labtech) using monochromator settings of 469-497 nm excitation / 500-560 nm emission. Cooperativity analyses were performed by nonlinear least-squares fitting to the Hill equation^50^: 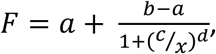 where *F* is the fluorescence signal, *x* is the concentration of RNA, and *a, b, c*, and *d* are fit parameters such that is the Hill coefficient.

## Results

We first report the characteristics of the genome exposure assay under standardized conditions of RNA-dye binding in the absence of membrane fusion. We then present results for influenza viral genome exposure during fusion, compare to a previously reported content-mixing probe, and test the dependence of fusion kinetics on target membrane curvature.

DiYO-1 (CAS 143413-85-8, marketed by Invitrogen as YOYO-1 and referred to here as DiYO) is a dimeric carbocyanine dye that is well described as intercalating into double-stranded DNA or single-stranded DNA and undergoing an approximately 1000-fold increase in fluorescence quantum yield^51-54^. Here, we show that it also binds with high affinity to single-stranded RNA (Fig. 2). The Hill coefficient for binding is approximately 2, consistent with cooperative dimeric binding and intercalation. Bulk fluorescence experiments (Fig. 2b) were used to select a dye concentration of 10 µM for liposomal encapsulation. Bulk fluorescence at any dye concentration was negligible in the absence of RNA.

**Figure 2.**
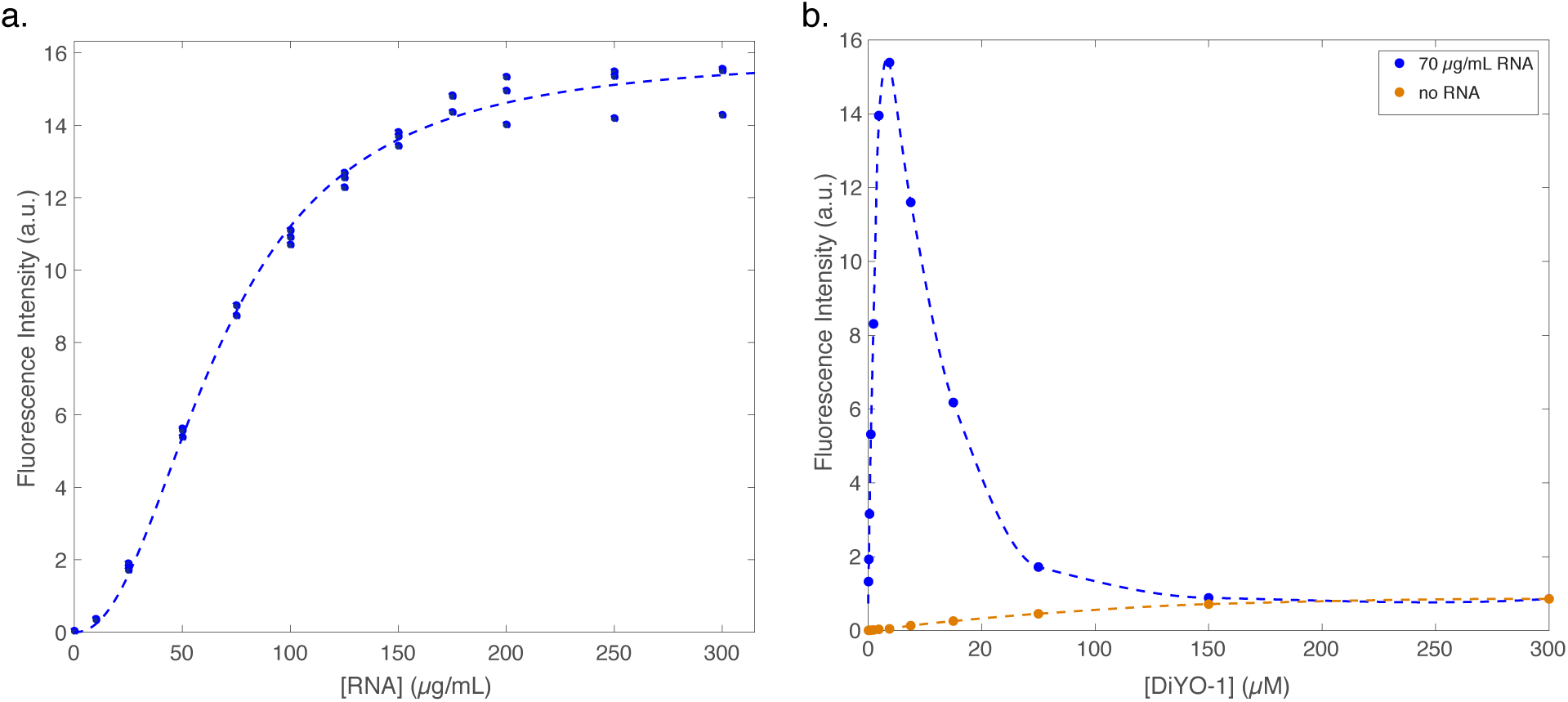
Dose-response of DiYO fluorescence upon incubation with RNA. A 10 µM solution of DiYO dye was incubated with increasing quantities of poly-A RNA to obtain a dose-response curve for DiYO fluorescence. Data (measured in triplicate) are plotted in panel (a) along with a fitted Hill equation curve. The best-fit Hill coefficient was 2.2 +/-0.3, and the EC_50_ was 67 µg/mL NA. The maximal fluorescence increase observed was 340-fold over the lowest amount of RNA tested. Plotted in panel (b) is the dose-response with a fixed amount of poly-A RNA and variable DiYO concentration.

Liposomes (67:20:10:2:1 POPC:DOPE:Chol:GD1a:DPPE-biotin) were extruded at 100 or 200 nm as indicated and immobilized in a microfluidic flow cell as previously described^47^. When the pH was decreased from 7.4 to 5.0 either in the exterior environment via simple buffer exchange or intraluminally via buffer exchange in the presence of FCCP ionophore, no change in background fluorescence was observed (Fig. 3). However, a robust increase in fluorescence was observed upon acidification after co-incubation with influenza virus (Fig. 1) although not before. This is consistent with detection of influenza genome exposure via a fusion pore.

**Figure 3.**
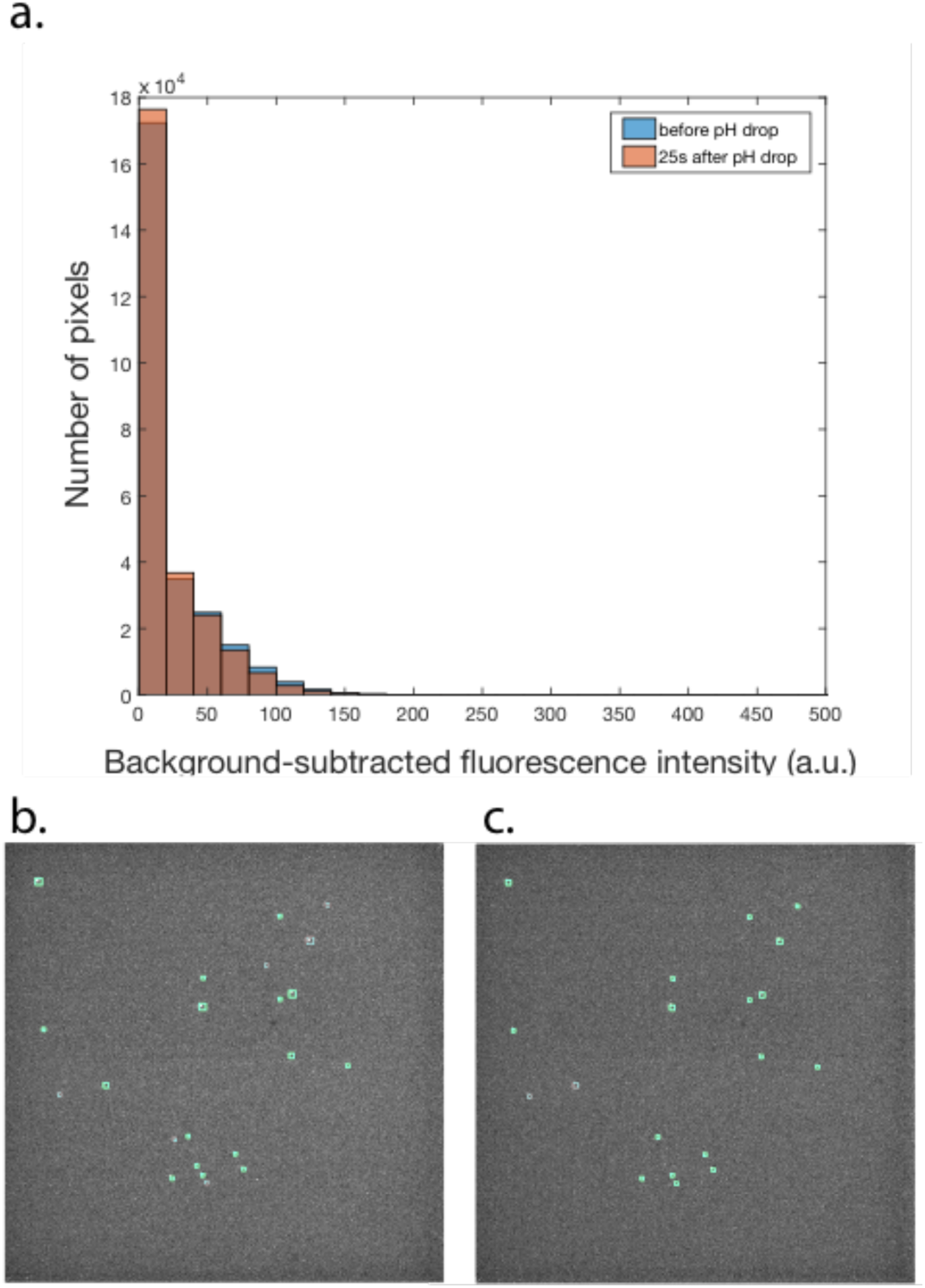
Encapsulated DiYO does not show measurable fluorescence gain in the absence of virus. Single-liposome DiYO intensities were measured at pH 7.0 and after intraluminal pH exchange to pH 5.0 using acidic buffer in the presence of ionophore FCCP. Overlaid intensity histograms are rendered in panel (a) and before/after images in panels (b) and (c), respectively. No significant fluorescence intensity gain was observed for the imaged liposomes in the absence of influenza virus (p > 0.8, Kolmogorov-Smirnov test). Green circles indicate single-liposome spots.

Single-event time-courses for genome exposure were measured for >100 influenza virus-liposome conjugates per condition and compiled into cumulative distribution functions (CDFs; Fig. 4). These are compared to CDFs measured under identical conditions for lipid dye mixing between the virus and liposome. For both 100- and 200-nm liposomes, genome exposure occurred more slowly than lipid mixing, but the time-courses were not statistically distinguishable between liposome sizes (p > 0.5, Kolmogorov-Smirnov test). Single-event analysis also indicated that genome exposure requires more kinetic steps contributing to the rate-limiting processes than lipid mixing (Table S1). This is as expected given prior theory ^1, 55^ of the free-energy barriers to fusion pore formation and any potential additional barriers to genome exposure, since genome exposure would require all the steps required for lipid mixing, then all those required for fusion pore formation, and possibly additional ones. If genome exposure or fusion pore formation were substantially slower than lipid mixing, those processes could dominate the rate-limiting step, but that does not appear to be the case.

**Figure 4.**
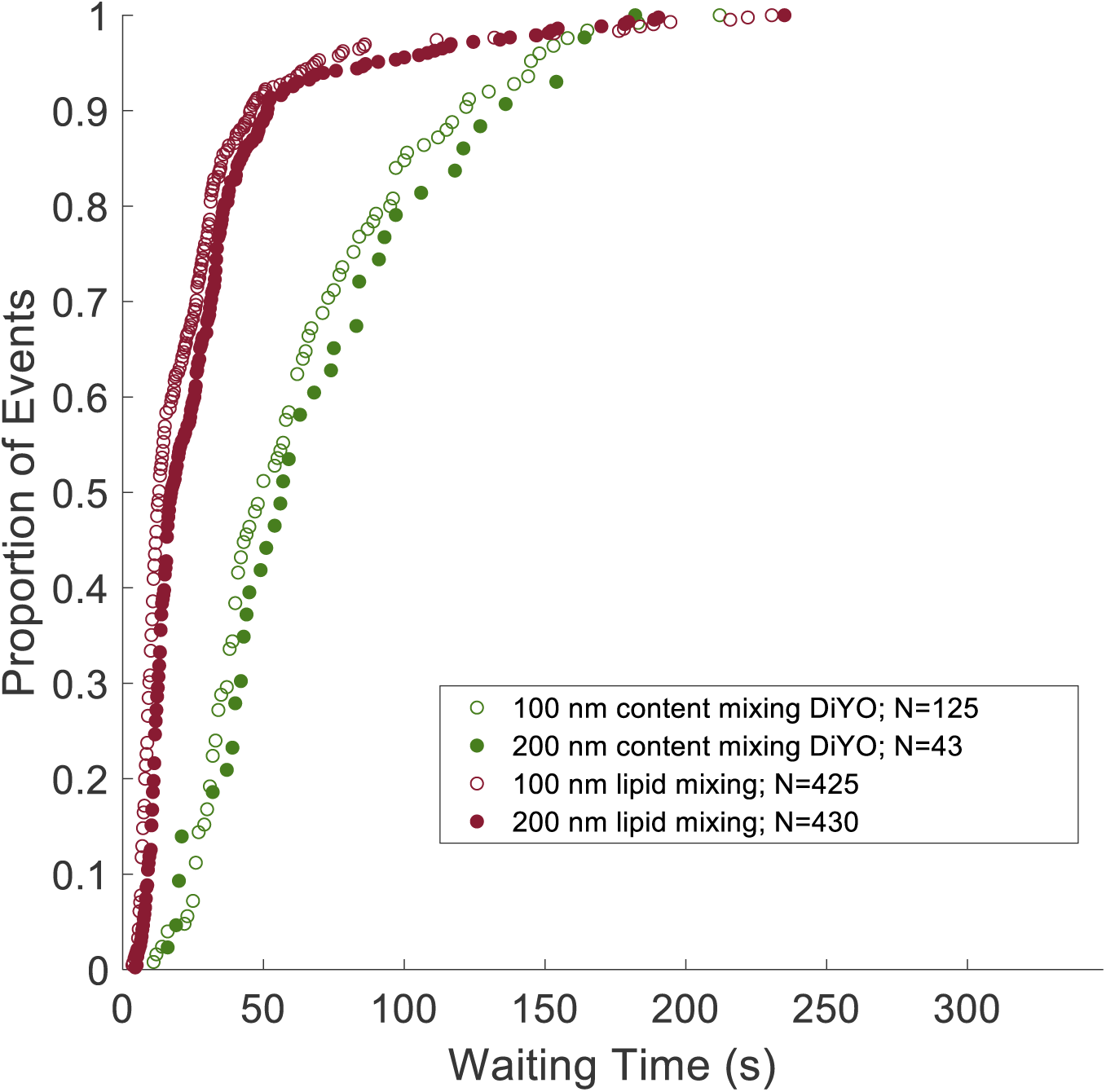
Genome exposure occurs more slowly than lipid mixing but with the rate-limiting step similarly independent of target membrane curvature. Influenza virus was bound to 100-nm or 200-nm target liposomes, fusion triggered via a drop in pH, and single-event kinetics of either lipid mixing or genome exposure monitored via fluorescence microscopy. No major differences in kinetics were observed between target liposome populations. As expected, genome exposure occurred more slowly than lipid mixing, and not all lipid mixing events led to genome exposure. These observations are consistent with one or more additional free-energy barriers for fusion pore formation and potentially downstream events. Fit parameters and single-event statistics are given in Table S1.

For comparison, we performed a calcein content-mixing assay where target liposomes were loaded with calcein dye instead of DiYO and analogous single-particle fluorescence traces were measured. Similar to previous observations in non-viral systems^38, 56-58^, we observed several patterns of fluorescence traces, which we interpret as corresponding to liposome bursting, leakage of liposome contents, and fusion. Under these conditions, the calcein assay does not differentiate leakage and fusion. Characteristic traces are shown in Figure 5. Under the labeling and illumination conditions used, no illumination-dependent fusion was observed as had been reported previously for SNARE-based fusion^38^. If the bursting traces are discarded, then aggregate slow leakage and fusion traces can be assembled into a cumulative distribution function (Fig. 5). The kinetics of calcein combined fusion and leakage events (“fusion+leakage”) are somewhat slower than those of genome exposure, potentially suggesting that late leakage events may occur in virus-liposome conjugates without forming productive fusion pores. These results illustrate the selectivity of the DiYO genome exposure assay for productive fusion events, since an increase in DiYO fluorescence requires intraluminal mixing of dye and RNA.

**Figure 5.**
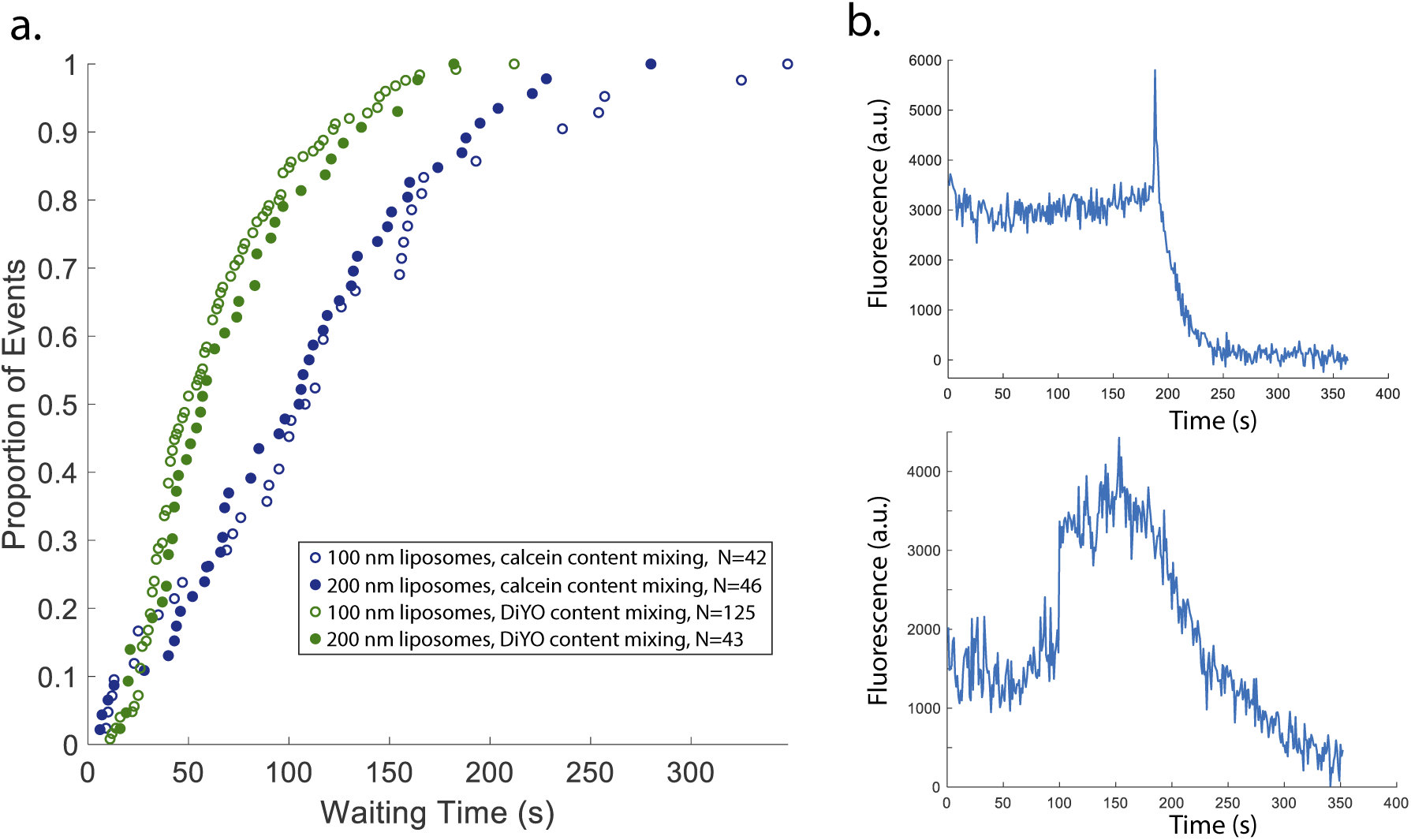
Calcein dequenching and DiYO RNA binding both show kinetics that do not depend on liposome curvature. Cumulative distribution functions for calcein dequenching and DiYO fluorescence are shown in panel (a); both display indistinguishable kinetics when measured with 100-nm and 200-nm liposomes. Representative single-virus calcein dequenching traces are shown in panel (b). Two profiles are observed: one (top panel) is attributed to liposome rupture, and the other (bottom panel) is attributed to either liposome leakage or fusion; a mixture of such events likely exists in the population.

In these experiments, we detected 28-40 lipid mixing events (mean of 35) per 100 Texas-Red-labeled viral particles (defined as Texas-Red-labeled and immunopositive for hemagglutinin) and 4-20 DiYO genome exposure events (mean of 16) per 100. Compared across biological replicas, there was no significant difference in DiYO fluorescence events versus calcein fusion+leakage events (p > 0.7, Kolmogorov-Smirnov test) and no significant difference in event frequencies between 100-nm and 200-nm particles (p > 0.4, Kolmogorov-Smirnov test). The lower fraction of viral particles undergoing genome exposure is consistent with prior measurements of lipid versus content mixing in influenza virus^32^ and may also reflect physiological inefficiency in successful infection^59-61^.

## Discussion

We have presented the use of an RNA-binding dye to measure the kinetics of influenza genome exposure during viral entry. This assay permits single-virus measurements of a step in viral entry equivalent to or downstream of content mixing between an influenza virion and the content of a target membrane. RNA binding by DiYO is highly sensitive and specifically measures exposure of the viral genome to target membrane contents; target leakage and rupture events do not register in this assay. It thus provides an approach for detecting productive fusion events in viral entry.

Given the relative rapidity and lack of cellular factors required to detect DiYO-influenza RNA binding, we hypothesize that exposure events occur as follows. The M1 layer undergoes partial disassembly, likely prior to fusion itself, and dye then diffuses into the virion at the time of fusion pore formation, binding to exposed portions of the viral RNPs. It is of course possible that the viral genome instead diffuses into the lumen of the target liposome, but full disassembly of the influenza virion is thought to require additional factors^5, 62^ and is likely slower than the events detected here. It would be valuable to develop sensors for viral RNA that could be used for real-time single-virus detection and that are incapable of diffusing through fusion pores. Such probes could be used as additional measures of downstream events in entry of influenza and other enveloped RNA viruses. Aptamer approaches have indeed been used for sensitive detection of viral nucleic acid but have not yet been deployed with single-virus sensitivity^35, 36^.

Using this approach, we compare the kinetics of influenza genome exposure when the virus fuses to target liposomes of differing size and curvature. In contrast to prior reports on SNARE-mediated fusion^48^, we observe identical genome exposure kinetics with 100-nm and 200-nm target liposomes. In prior work, we demonstrated that influenza virus undergoes lipid mixing at indistinguishable rates to target liposomes of differing size^63^ but that membrane deformability is a controlling factor for lipid mixing^47^. It is thus not surprising that genome exposure, which occurs no earlier than content mixing and definitely downstream of lipid mixing, would show a similar independence of starting curvature. Classical membrane curvature theory^64^ dictates that smaller vesicles should have a more favorable free energy of fusion with influenza virions than larger vesicles because of the curvature stress thus relieved. Prior theoretical results on protein-free membrane fusion^65-68^ are largely in accordance with this but focus on the curvature energy of highly curved stalk intermediates. The results presented here demonstrate that the starting curvature does not measurably contribute to the rate limiting steps of either lipid or content mixing in influenza entry. This stands in contrast to the prior results for SNARE-mediated fusion^48^. It is of course possible that technical details of the assays and probes used could account for the difference, but the more interesting possibility is that this difference in curvature dependence illuminates the mechanistic differences between SNARE-mediated and viral membrane fusion.

In summary, nucleic acid binding by DiYO-1 yields a highly sensitive and specific assay for exposure of RNA virus genomes. We have leveraged this as a way to measure viral membrane fusion in real time with single-virion resolution, demonstrated here for influenza. We anticipate that this assay will further aid in unraveling the complex mechanisms underpinning enveloped virus entry.

## Author Contributions

A.M.V.G. designed research, performed experiments, analyzed data, and wrote the manuscript. S.M. performed experiments and analyzed data. P.M.K. designed research, analyzed data, and wrote the manuscript.

## Declaration of Interests

The authors declare no competing interests.

## Acknowledgements

The authors thank S. Boxer for many helpful conversations. This work was supported by a Wallenberg Academy Fellowship to P.M.K. and by the Swedish Research Council (2017-04236 to P.M.K.).

## Notes

### Competing Interest Statement

The authors have declared no competing interest.

