## Supplementary material for "Measuring single-virus fusion kinetics using an assay for nucleic acid exposure": Table S1

| **Fit Parameter** | **Tau (s)** | | **Lag time (s)** | | **N** | | **$N_{min}$** | |
|---|---|---|---|---|---|---|---|---|
| **Liposome size** | 100 nm | 200 nm | 100 nm | 200 nm | 100 nm | 200 nm | 100 nm | 200 nm |
| Lipid mixing (Texas Red dequenching) | 22 (19-30) | 23 (19-27) | 4.7 (4.2-5.5) | 5.5 (5.0-6.4) | 0.78 (0.56-0.91) | 0.95 (0.78-1.1) | 0.6 (0.5-0.9) | 0.9 (0.7-1.1) |
| Genome exposure (DiYO dequenching) | 30 (22-38) | 34 (18-56) | 9.7 (5.6-13) | 9.4 ($10^{-4}$-16) | 1.8 (1.4-2.4) | 1.8 (1.0-3.6) | 2.1 (2.1-3.1) | 2.9 (2.1-3.4) |
| Content mixing + leakage (calcein) | 158 (100-250) | 69 (34-141) | 6 (4.9-12) | $4.7\times10^{-6}$ ($2.2\times10^{-6}$ -25) | 0.62 (0.41-0.86) | 1.3 (0.50-2.0) | 1.0 (0.76-1.4) | 1.6 (1.1-2.8) |

**Table S1**. Kinetic fit parameters for lipid and content dyes. Maximum-likelihood parameters are listed from fitting to a lagged gamma CDF: $P(x \mid N, \tau, g) = \frac{1}{\tau^N \Gamma(N)} \int_0^{x-g} t^{N-1} e^{-t/\tau} dt$, where $N$, $\tau$, and $g$ are shape, scale, and lag respectively. $N_{min}$ is calculated as the inverse of the randomness parameter: $r \equiv \langle (t - \langle t \rangle)^2 \rangle / \langle t \rangle^2$ for single-event waiting times t[1, 2]. 90% confidence intervals are given in parentheses. Lipid mixing occurs most rapidly; genome exposure occurs following that and in a multi-step kinetic fashion. Interestingly, content mixing+leakage occurs in a fashion dominated by a single-exponential process ($N_{min}$ ~ 1).
